# Exploring the metastatic role of the inhibitor of apoptosis BIRC6 in Breast Cancer

**DOI:** 10.1101/2021.04.08.438518

**Authors:** Santiago M. Gómez Bergna, Abril Marchesini, Leslie C. Amorós Morales, Paula N. Arrías, Hernán G. Farina, Víctor Romanowski, M. Florencia Gottardo, Matias L. Pidre

## Abstract

Breast cancer is the most common cancer as well as the first cause of death by cancer in women worldwide. BIRC6 (baculoviral IAP repeat-containing protein 6) is a member of the inhibitors of apoptosis protein family thought to play an important role in the progression or chemoresistance of many cancers. The aim of the present work was to investigate the role of apoptosis inhibitor BIRC6 in breast cancer, focusing particularly on its involvement in the metastatic cascade.

We analyzed BIRC6 mRNA expression levels and Copy Number Variations (CNV) in three breast cancer databases from The Cancer Genome Atlas (TCGA) comparing clinical and molecular attributes. Genomic analysis was performed using CBioportal platform while transcriptomic studies (mRNA expression levels, correlation heatmaps, survival plots and Gene Ontology) were performed with USC Xena and R. Statistical significance was set at p-values less than 0.05.

Our analyses showed that there was a differential expression of BIRC6 in cancer samples when compared to normal samples. CNV that involve amplification and gain of BIRC6 gene were correlated with negative hormone receptor tumors, higher prognostic indexes, younger age at diagnosis and both chemotherapy and radiotherapy administration. Transcriptomic and gene-ontology analyses showed that, in conditions of high BIRC6 mRNA levels, there are differential expression patterns in apoptotic, proliferation, and metastatic pathways.

In summary, our *in silico* analyses suggest that BIRC6 exhibits an antiapoptotic, pro-proliferative and an apparent pro-metastatic role and could be a relevant molecular target for treatment of Breast Cancer tumors.

## 1. Introduction

Breast cancer (BC) is one of the most prevalent cancers in the general population, and it’s the leading cause of death for female cancer patients. This pathology presents heterogeneity in the biological behavior of tumors and a great clinical variability (1). The main cause of breast cancer-related death is the development of metastasis, which accounts for 90% of deaths. The recurrence of this disease originates in local processes on secondary organs and is associated with a poor prognosis (2). The main characteristic that differentiates a benign tumor from a malignant tumor is the invasiveness of the malignant cells. This ability to invade surrounding tissues is fundamental for metastasis. Said process comprises several steps, all of which are controlled by different cellular and environmental signals (3). Understanding metastasis is of central importance when searching for antitumor therapies since identification of potential molecular targets for treatment requires discerning the key factors involved in it. Apoptosis is a highly regulated process and its failure can result in many pathological conditions including tumor development. In mammals, programmed cell death is usually regulated by the IAP family of proteins (named after their main function, IAP: “ inhibitors of apoptosis proteins”) (4). IAP, which were originally isolated and characterized from baculoviral genomes, contain highly conserved protein-protein interaction motifs called baculoviral IAP repeats (BIR). Through these BIR motifs, IAP are able to associate with different caspases and prevent their activity, thereby inhibiting apoptosis. Many IAPs also contain a RING domain at their C-terminal end with E3 ubiquitin ligase activity that allows control of protein levels by ubiquitination and degradation via proteasome (5). This family of proteins plays a central role in the control of survival and programmed cell death by regulating determining factors in both the caspase activation pathway as well as the NF-κB pathway (5).

The role of IAPs in tumor progression and metastasis has been reported for several tumor types (6–9). Recently, a PanCancer transcriptomic analysis showed a key role for IAPs in tumor physiology (10).

BIRC6 (Baculoviral IAP repeat-containing 6, also known as Apollon or BRUCE in mice) is a IAP of approximately 530 kDa that contains a BIR domain at its N-terminal region and an ubiquitin ligase domain at its C-terminal (UBC). Different research studies have described that BIRC6 plays a dual role as an anti-apoptotic IAP and as a chimeric E2/E3 ubiquitin ligase. BIRC6 is capable of catalyzing ubiquitination of different target proteins such as SMAC/DIABLO and Caspase-9, among others (11–13). BIRC6 not only inhibits the pro-apoptotic protein SMAC, but also binds to procaspase-9 and prevents its cleavage (13,14). Likewise, through its BIR domain it can bind and inhibit active caspases, including caspases 3, 6, 7 and 9 (13–16).

In addition to its function as an inhibitory protein of apoptosis, BIRC6 plays both an important role in cell proliferation and as a regulator of cytokinesis (17). BIRC6 is associated with the membrane and is located in the Golgi compartments and in the vesicular system (16). BIRC6 also participates in other cellular processes such as autophagy, in which it regulates autophagosome-lysosome fusion (18,19).

Different groups have demonstrated that IAP are overexpressed in several types of tumor cells, and it has been inferred that they could be related to tumorigenesis, treatment resistance, worse prognosis and oncogenesis (5,7,20–30). In particular, BIRC6 overexpression has been found in tumor tissues of gastric carcinomas (31), colorectal cancer (21), breast cancer (22), and lung cancer (24) among others. These findings postulate IAP, and particularly BIRC6, as a potential therapeutic target against different cancers, especially those that most frequently develop chemoresistance. Such is the case of BC, for which BIRC6 has not yet been completely validated as a therapeutic target. Our aim was to further evaluate whether BIRC6 may play a role in BC and metastasis using bioinformatic tools.

## 2. Materials and methods

### 2.1 Breast Cancer and normal tissue samples

We used different public datasets containing clinical, genomic and transcriptomic information from patient samples. We also used two different platforms to analyze the data.

For these analyses, the TCGA and GTEx databases and the UCSCXena platform were employed. Samples corresponding to mammary tissues were filtered and the expression of BIRC6 and other genes was evaluated in conditions of normal tissue and tissue of primary tumors.

### 2.2 CNV and clinical attributes

For these analyses, the BC database METABRIC (32,33) and the cBioPortal platform were used. The BIRC6 gene was used as a query and the correlation of different copy number variations (CNV) with the expression of BIRC6 and several clinical attributes of interest was assessed.

### 2.3 Survival plots

The UCSCXena (34) platform and the TCGA Pan-Cancer (35–38) database were used to analyze the patients survival. To this end, samples corresponding to BC were filtered and the expression of BIRC6 were evaluated. In the analysis, the samples were divided into two groups: high (≥11.1) and low expression of BIRC6 (<11.1), and survival rates were plotted.

### 2.4 Transcriptomic analyses

The UCSCXena (34) and cBioPortal (39,40) platforms and the TCGA GTEx, TCGA Pan-Cancer (35–38) and Molecular Taxonomy of BC International Consortium (BC, METABRIC) databases were used to evaluate the expression of BIRC6 (32,33). TCGA GTEx was selected because it is the only one that includes transcriptomic data from normal tissue from healthy volunteers and tissue from primary tumors obtained from patients with BC. The TCGA Pan-Cancer (PANCAN) and BC (METABRIC (32,33)) databases were selected on the basis of the number of samples included in the datasets, and the different parameters that could be evaluated. In the case of TCGA Pan-Cancer and TCGA-Target-GTEx, normalized RNAseqV2 data was employed using RSEM quantification (41). In the case of BC (METABRIC), the mRNA expression data in z-scores relative to all samples (log microarray) carried out on the Illumina HT-12 v3 platform (Illumina_Human_WG-v3) (32) was used.

For different pathway correlation analyses, the TCGA Pan-Cancer database (PANCAN) and the UCSCXena and cBioPortal platforms were used. Samples corresponding to BC were filtered and the expression of BIRC6 and different molecules involved in the metastatic cascade pathways were evaluated. In the analysis, the samples were divided into two groups: high expression of BIRC6 (≥11.1) and low expression of BIRC6 (<11.1). In addition, the analysis was performed for different types of BC: hormone receptor positive and hormone receptor negative. The cut-off points to consider HR positive were the following: ESR1 ≥ 10 and PGR ≥7 for RSEM normalized expression.

### 2.5 Gene Ontology and pathway analysis

Publicly available data corresponding to the TCGA-BC dataset was used to perform differential gene expression analysis and Gene Ontology. BC harmonized data (hg38) in HTseq-Counts (raw counts) format was downloaded from the TCGA database (https://portal.gdc.cancer.gov/) using the GDCdownload function of the TCGABiolinks package (2.18.0) (42–44) in R (45). The dataset contained raw, fully sequenced transcriptome data from 1,088 primary tumor samples from BC patients.

To perform the differential expression analysis of genes, the samples were separated into two groups: those that presented BIRC6 expression values greater than the median and those that presented BIRC6 expression values lower than the median. All samples were normalized and filtered using R / Bioconductor’s TCGABiolinks package following the standard pipeline. They were preprocessed using the TCGAanalyze_Preprocessing function and a correlation cutoff of 0.6, then normalized with TCGAanalyze_Normalization using the GC content method, and finally filtered using TCGAanalyze_Filtering by quantile as recommended.

For the enrichment analysis, the TCGAanalyze_EAcomplete function was applied using the DEGs (Differential Expressed Gene) with a log(FC)>0 for overexpressed DEGs or log(FC)<0 for less expressed DEGs, in order to obtain the 3 ontologies of those genes, respectively (GO: biological process, GO: cellular component, and GO: molecular function) and the pathways in which they were involved. These results were plotted using the TCGAvisualize_EAbarplot function, showing the 35 biological processes with the lowest FDR.

### 2.6 Statistical analysis

*Expression of BIRC6 in samples of healthy volunteers and samples of mammary tumor tissue*: n = 1275 (n = 1099 primary tumors; n = 176 normal breast tissue); t-Test. *Expression of BIRC6 vs. copy number*: n = 2173 (n = 232 Deletions; n = 1818 Diploids; n = 111 Gains; n = 12 Amplifications); multiple ANOVA followed by TukeyHSD. *BIRC6 CNV vs. presence / absence of receptors*: n = 2140 was used to evaluate the estrogen receptor (n = 1617 ER +; n = 523 ER-), n = 1980 to evaluate the progesterone receptor (n = 1040 PR +; n = 940 PR -) and n = 1980 to evaluate the HER2 receptor (n = 247 HER2 +; n = 1733 HER2 -); χ2 test grouping CNV that implied an increase in the number of copies (gains and amplifications) and those that did not imply it (deletions and diploids). *BIRC6 CNV vs. age at diagnosis and the Nottingham Prognostic Index*: n = 2173 (n = 232 Deletions; n = 1818 Diploids; n = 111 Gains; n = 12 Amplifications); multiple ANOVA followed by TukeyHSD. *BIRC6 CNV vs. Neoplastic Histological Grade*: n = 2072 (n = 174 Grade 1; n = 851 Grade 2; n = 1047 Grade 3); χ2 test grouping CNV that implied an increase in the number of copies of the gene (gains and amplifications) and those that did not imply it (deletions and diploids). *BIRC6 CNV vs. treatment with chemotherapy or radiotherapy*: n = 1980 (n = 1173 Treated; n = 807 Not Treated) in the case of radiotherapy and n = 1980 (n = 411 Treated; n = 1569 Untreated) in the case of chemotherapy; χ2 test by grouping those CNV that implied an increase in the number of copies (gains and amplifications) and those that did not imply it (deletions and diploids). *BIRC6 expression vs. proteins of different pathways*: n = 1211 (n = 697 High Expression of BIRC6; n = 514 Low Expression of BIRC6); t-test. *Correlation analysis*: Pearson’s correlation coefficient. Statistical significance was considered when p-value did not exceed 0.05 for all studies.

## 3. Results

### 3.1 Patient cohort

For clinical, genomic and transcriptomic analyses three different databases were employed. Patient attributes of each database are summarized in Table 1. The numbers in each cell indicate the number of patients with the corresponding attribute.

**Table 1.** Patient cohort. Table 1 shows principal clinical attributes corresponding to each of three databases used in this work. The number in each cell indicates the quantity of patients with clinical attributes specified in column one.

| Clinical attribute |  | Database |  |  |
| --- | --- | --- | --- | --- |
|  |  | METABRIC | TCGA, PanCancer | TCGA-BRCA 2021 |
| Number of patients (BC) |  | 2509 | 1084 | 1098 |
| Age | ≤ 50 | 567 | 294 | 297 |
|  | > 50 | 1926 | 756 | 766 |
| Histological Grade | 1 | 214 | Not available | Not available |
|  | 2 | 976 | Not available | Not available |
|  | 3 | 1198 | Not available | Not available |
| ER Status | Positive | 1825 | Not available | Not available |
|  | Negative | 644 | Not available | Not available |
| PR Status | Positive | 1040 | Not available | Not available |
|  | Negative | 940 | Not available | Not available |
| HER2 Status | Positive | 247 | Not available | Not available |
|  | Negative | 1733 | Not available | Not available |
| Hormone Therapy | Yes | 1216 | Not available | Not available |
|  | No | 764 | Not available | Not available |
| Chemotherapy | Yes | 412 | Not available | 1097 |
|  | No | 1568 | Not available | 1097 |
| Radiotherapy | Yes | 1173 | 549 | 1097 |
|  | No | 807 | 434 | 1097 |

### 3.2 BIRC6 is differentially expressed in tumor samples

In order to evaluate the role of BIRC6 in human BC samples, we proceeded to study transcriptomic databases.

The expression of BIRC6 was compared in samples from primary tumors of BC patients and normal tissue samples from healthy volunteers. The result is shown in Fig 1A. We observed a statistically significant increase in the expression of BIRC6 in primary tumor samples in comparison to normal tissue from healthy volunteers. Cancer patients had a median z-score of 0.113, whilst normal tissue has a median z-score of -0.372.

**Fig 1.**
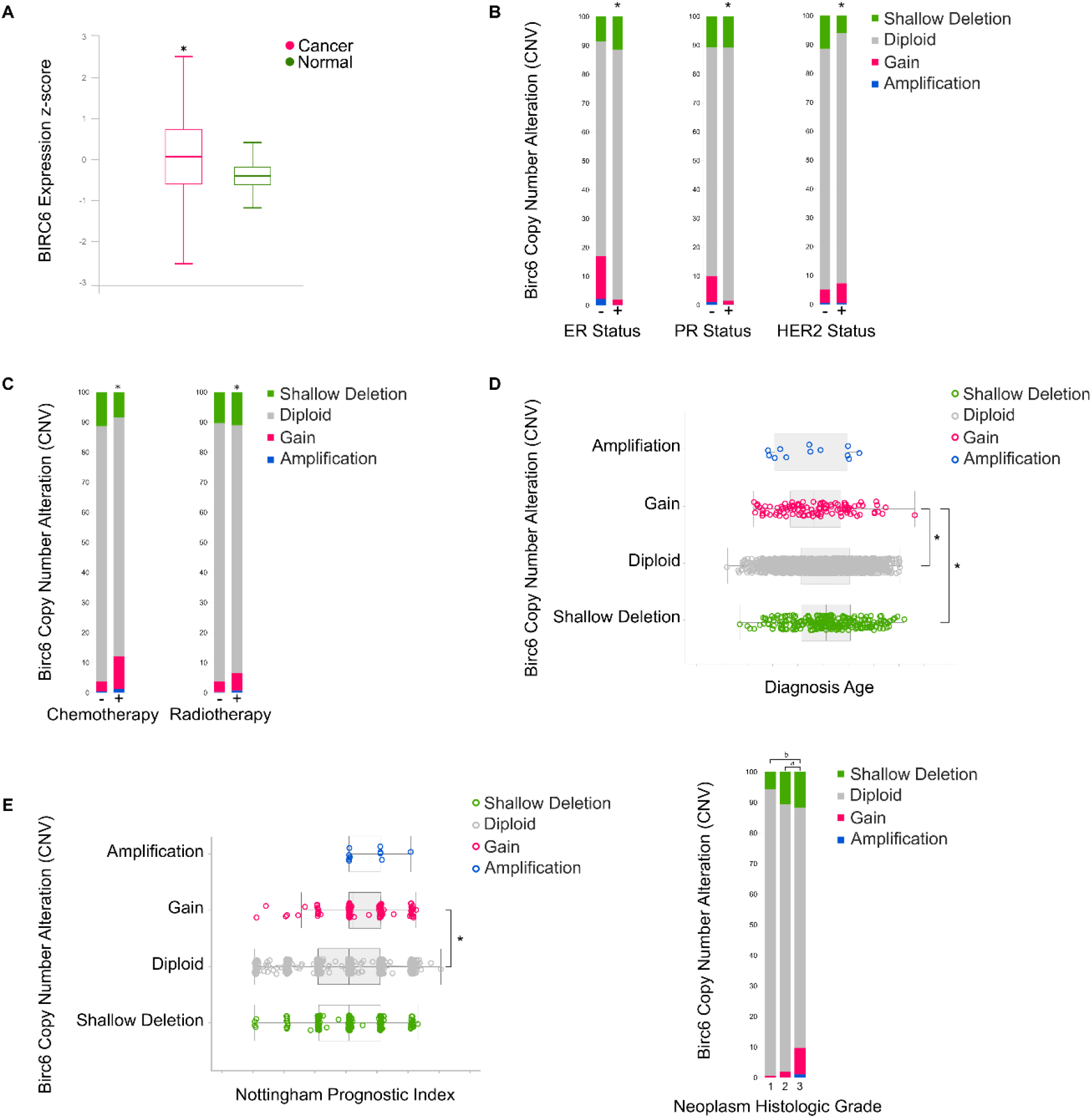
BIRC6 expression and copy number variation (CNV). (A) Differential expression of BIRC6 between tumor and normal tissue. (B) BIRC6 CNV abundance vs. different clinical attributes: ER, PR and HER2 status, χ2 *p<0.05; (C) chemotherapy and radiotherapy, χ2 *p<0.05; (D) diagnosis age, multiple ANOVA followed by TukeyHSD *p<0.05; (E) Notingham prognostic index, multiple ANOVA followed by TukeyHSD *p<0.05 and neoplasm histologic grade, χ2 *p<0.05.

### 3.3 Increased copy number of the BIRC6 gene correlates with higher cellular dedifferentiation and worse prognosis

To characterize whether the alteration of BIRC6 has an impact on the different clinical attributes of patients we first evaluated if BIRC6 gene copy number variation (CNV) implied any change of its expression level. The CNV data was divided into four groups: amplifications, gains, diploids and deletions. A direct relationship was found between copy number and mRNA expression. Samples with gene amplifications showed the highest expression levels whereas genomic deletions were correlated with samples exhibiting the lowest expression levels (S1 Fig).

Following these results, we decided to evaluate the relationship between CNV and the presence or absence of estrogen (ER), progesterone (PR) and epidermal growth hormone 2 (HER2) receptors. The absence of these receptors in BC is associated with a much more aggressive phenotype and an inability to use hormonal therapies to stop its development (Fig 1B). We observed a higher proportion of amplifications and gains of CNV in ER (-) and PR (-) samples than in ER (+) and PR (+) samples, respectively.

We evaluated BIRC6 CNV distribution according to chemotherapy and radiotherapy treatments. Fig 1C shows that in samples from patients who received either of the two therapies, there was a greater amplification and gain percentage compared to those who did not.

After evaluation of the average age at which patients with the different CNVs were diagnosed, it became apparent that patients with higher CNVs tended to be diagnosed at a younger age than those who maintained diploidy or had deletions in BIRC6 gene (Fig 1D). We assessed the influence of CNV on two parameters that reflect prognosis: Nottingham Index and the Neoplastic Histological Grade. It was observed that patients with amplifications had a higher Nottingham Index than the rest of the conditions, thus implying worse prognosis (Fig 1E). Furthermore, we found a higher proportion of patients with amplifications and gains of the BIRC6 gene in patient samples with Histological Grade 3 (Fig 1E).

### 3.4 BIRC6 expression and survival

Breast cancer patient survival was evaluated using data of 1084 samples from TCGA PanCancer Atlas. Patients were divided into two groups, high and low BIRC6 expression, and survival was plotted for both groups. Survival time was significantly lower for patients with higher BIRC6 expression levels (Fig 2A).

**Fig 2.**
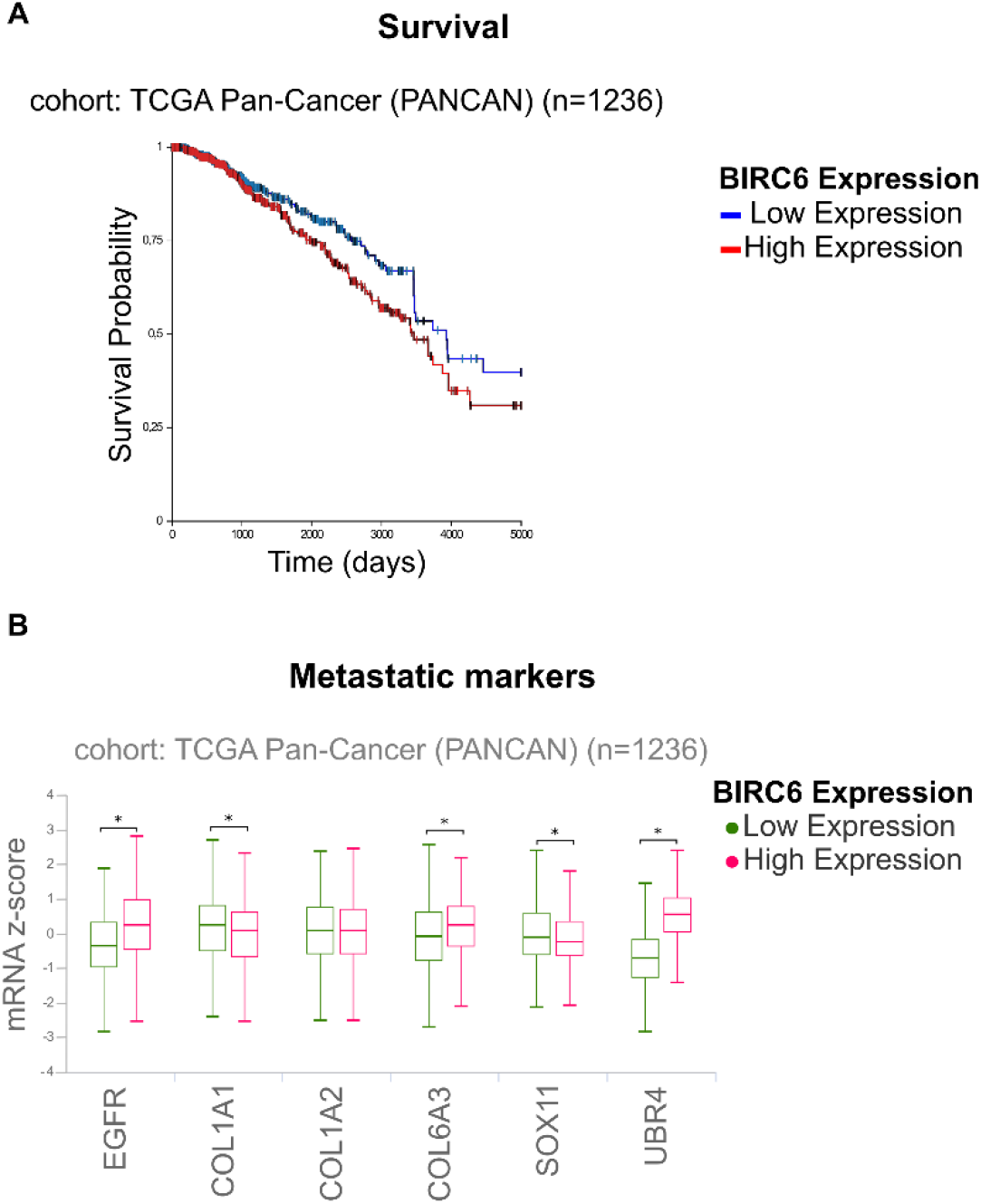
Survival and metastatic markers. (A) Survival plot. BC samples were filtered from TCGA Pan-Cancer and the expression of BIRC6 was evaluated. In the analysis, the samples were divided into two groups: high expression of BIRC6 and low expression of BIRC6 and survival were plotted for each group. T-test * p-value<0.05. (B) Metastatic markers expression. Boxplot of the expression (z-score) of common metastatic markers in condition of high and low BIRC6 expression. T-test * p-value<0.01.

### 3.5 BIRC6 expression and pathways

Differential expression of six general metastasis markers (EGFR, COL1A1, COL1A2, COL6A3, SOX11 and UBR4) was determined in patients with low and high expression of BIRC6. EGFR, COL6A3 and UBR4 showed a significant increase in their expression levels in samples with high BIRC6 expression (Fig 2B). Furthermore, we observed a significant increase in LDHA and a significant decrease in PDH mRNA levels in samples with high expression of BIRC6 in accordance with the expected metabolic switch for tumor cells (S2 Fig and S1 Table). For a more systematic approach we conducted a Differential Expression Analysis (DEA) followed by a gene ontology study. Differential genes obtained in this analysis were divided in two output groups: overexpressed (Fig 3) and less expressed (S3 Fig) genes, both under conditions of high BIRC6 expression levels. Fig 3 summarizes biological processes, cellular components, molecular function and pathways in which overexpressed genes are involved. We found that genes involved in gene expression (nucleosome and chromatin assembly and organization, protein-DNA complexes and DNA packaging) were overexpressed in the analyzed conditions as well as those involved in actin and myosin cytoskeleton signaling, epithelial adherence signaling and tight junction signaling.

**Fig 3.**
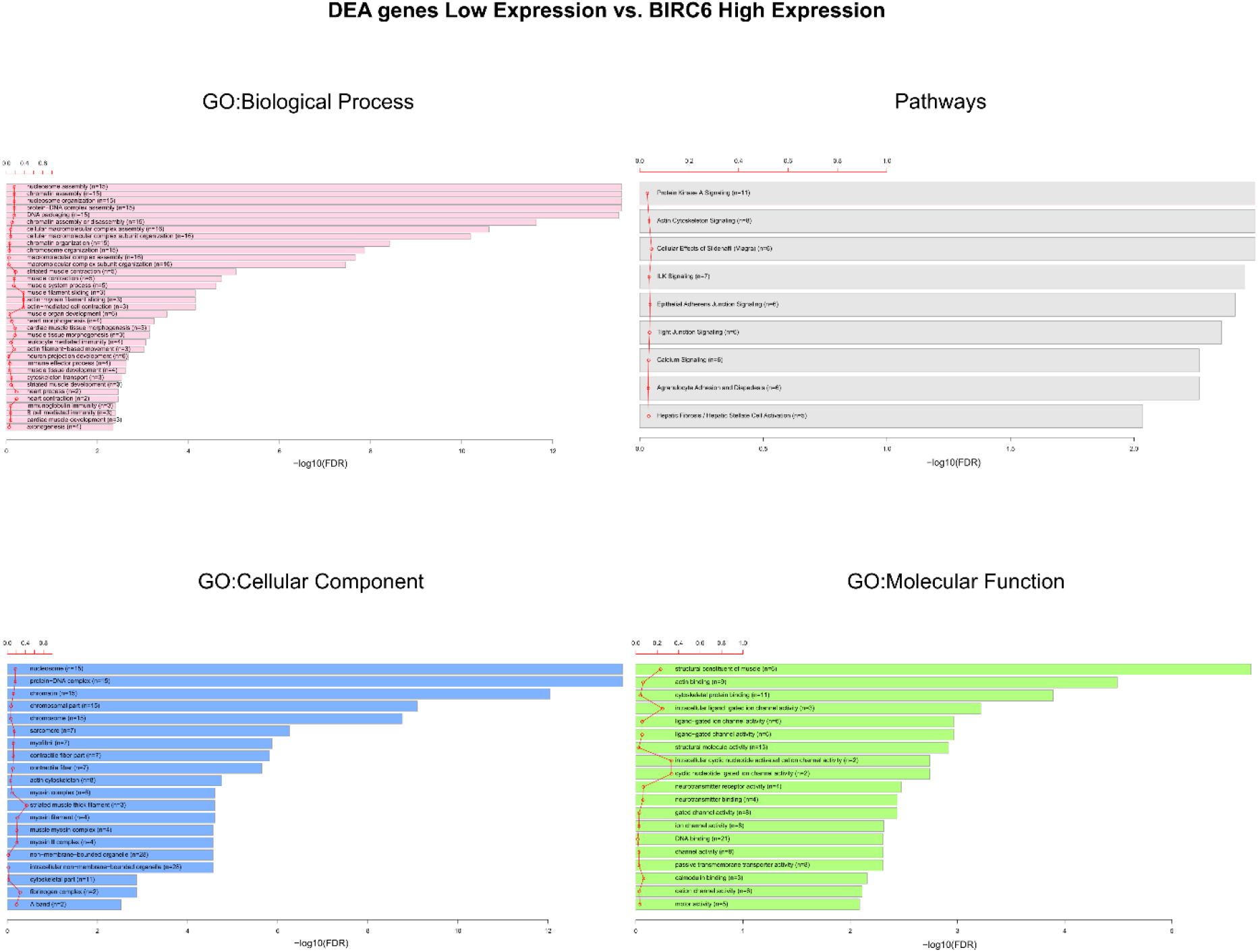
Gene ontology and over-represented pathways. Graphs show the canonical pathways significantly over-represented (enriched) by the DEGs (differentially expressed genes) with the number of genes for the main categories of the three ontologies (GO: biological process, GO: cellular component, and GO: molecular function, respectively). The statistically significant canonical pathways in the DEGs are listed according to their p-value corrected by FDR (-log10) (colored bars) and the ratio of the listed genes found in each pathway over the total number of genes in that pathway (ratio, red line).

These results suggest that BIRC6 could play an important role in tumor homeostasis and metastasis development. For this reason, we decided to perform a deeper analysis on apoptosis, proliferation, angiogenesis, migration and focal adhesion pathways.

#### i. Apoptosis

We evaluated the expression of Bax, Bcl-2, Caspase 9, Caspase 3, Caspase 8, TP53, Cytochrome C, NFκB1 and DIABLO under conditions of either low or high BIRC6 expression (Fig 4A). In addition, correlation between the expression of BIRC6 and the aforementioned proteins was analyzed (Fig 4B) by grouping samples according to four criteria: normal tissue, tumor tissue and positive or negative hormone receptor (HR + or HR-). The results showed that those samples with high expression of BIRC6 had lower expression of Bax, Caspase 9, Cytochrome C and DIABLO, and higher expression of Bcl-2, Caspase 3, Caspase 8 and NFκB1 when compared to samples with low BIRC6 expression. Furthermore, we observed that there was no statistically significant difference in TP53 expression levels between both groups (Fig 4B). Finally, we observed a differential correlation between samples of healthy tissue and tumor tissue.

**Fig 4.**
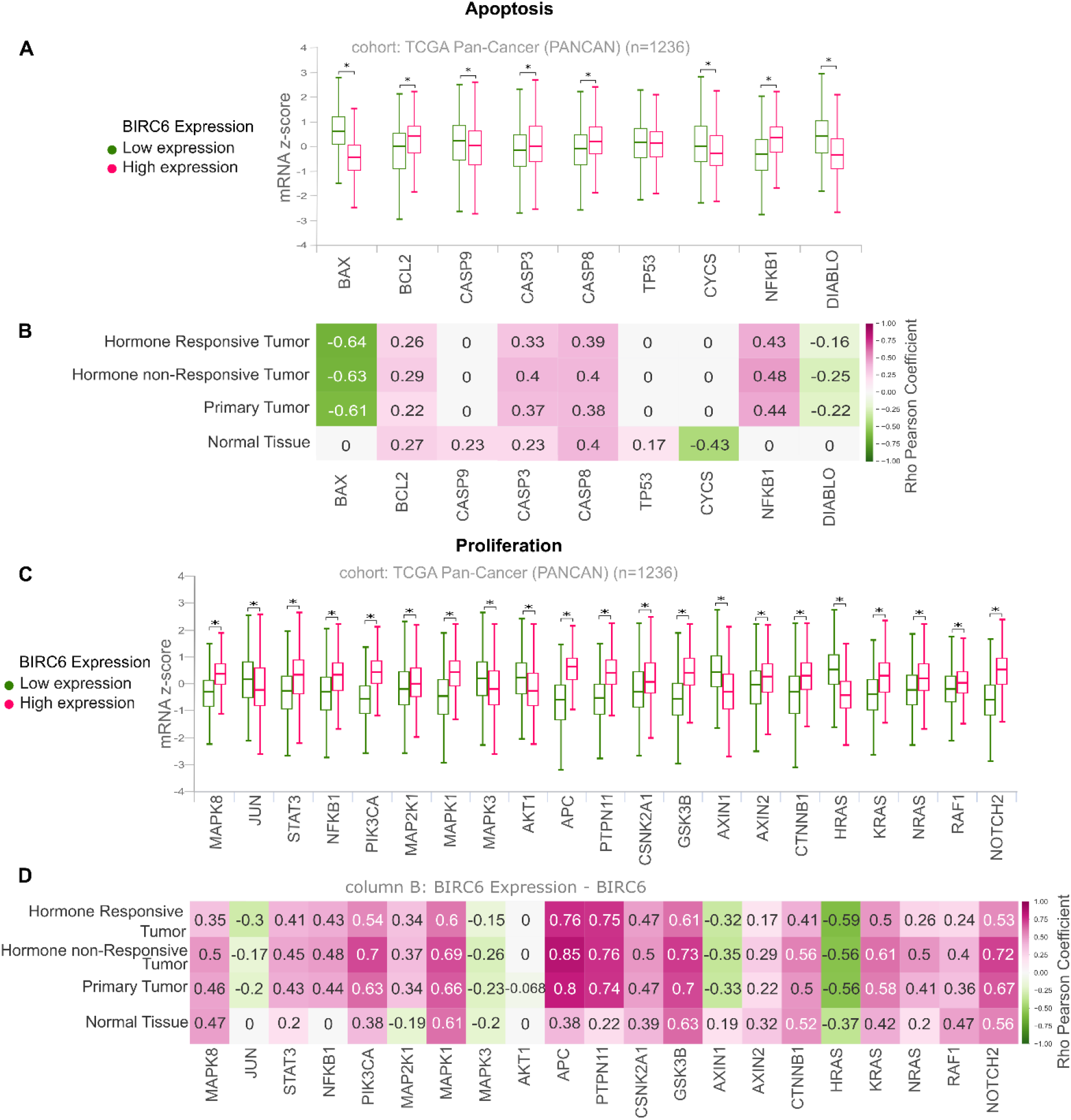
Transcriptomic analysis of apoptotic and proliferation pathways (TCGAPanCancer). (A) Boxplot Of The Expression (z-score) of proteins involved in the apoptotic pathway in condition of high and low BIRC6 expression. T-test *p-value<0.01. (B) Correlation between BIRC6 expression and proteins involved in the apoptotic pathway separated in: normal tissue, primary tumor, hormone responsive and hormone non responsive tumors. Numbers in cells indicate correlation pearson index in statistically significant comparisons. (C) Boxplot of the expression (z-score) of proteins involved in proliferative pathways in condition of high expression and low expression of BIRC6. T-test *p-value<0.01. (D) Correlation between BIRC6 expression and proteins involved in proliferative pathways separated in normal tissue, primary tumor, hormone responsive and hormone non responsive tumors. Numbers in cells indicate correlation pearson index in statistically significant comparisons.

#### ii. Proliferation

Another essential cellular process for tumor biology is proliferation, which is exacerbated in cancer cells. Taking this into account, we repeated the previous analysis using proteins linked to this process. We evaluated the expression of MAPK8, JUN, STAT3, NFkB1, PIK3CA, MAP2K1, MAPK1, MAPK3, AKT1, APC, PTPN11, CSNK2A1, GSK3B, AXIN1, AXIN2, CTNNB1, HRAS, KRAS, NRAS, RAF1 and NOTCH2 under conditions of low and high BIRC6 expression (Fig 4C). In addition, the correlation between the expression of these proteins and BIRC6 was analyzed in normal tissue, tumor tissue and positive or negative hormone receptors samples (Fig 4D). The results showed that those samples with high expression of BIRC6 have high expression of some proteins that promote proliferative pathways such as STAT3, PI3KCA, MAPKs and APC.

#### iii. Metastatic cascade: Angiogenesis, Migration and Focal Adhesion

The metastatic cascade consists of several steps: invasion, angiogenesis, surviving the passage through the circulatory system, adhesion and anchorage in distant organs and, finally, growth into micrometastases and, then, into consolidated metastases. We evaluated the correlation between BIRC6 overexpression and key regulators of the metastatic cascade. Angiogenesis is the generation of new blood vessels from pre-existing ones, which allow tumor cells to spread to distant organs. We assessed the expression levels of EGFR, IGF1R, INSR, HIF1A, ARNT, ANGPT, TEK and VEGFA in conditions of both low and high BIRC6 expression (Fig 5A). In addition, the correlation between the expression of these proteins and BIRC6 was analyzed in samples of normal tissue, tumor tissue and positive or negative hormone receptors (HR + or HR-). We found that those samples with high expression of BIRC6 have high expression levels of some angiogenesis promoters such as HIF1A and VEGFA (Fig 5B).

**Fig 5.**
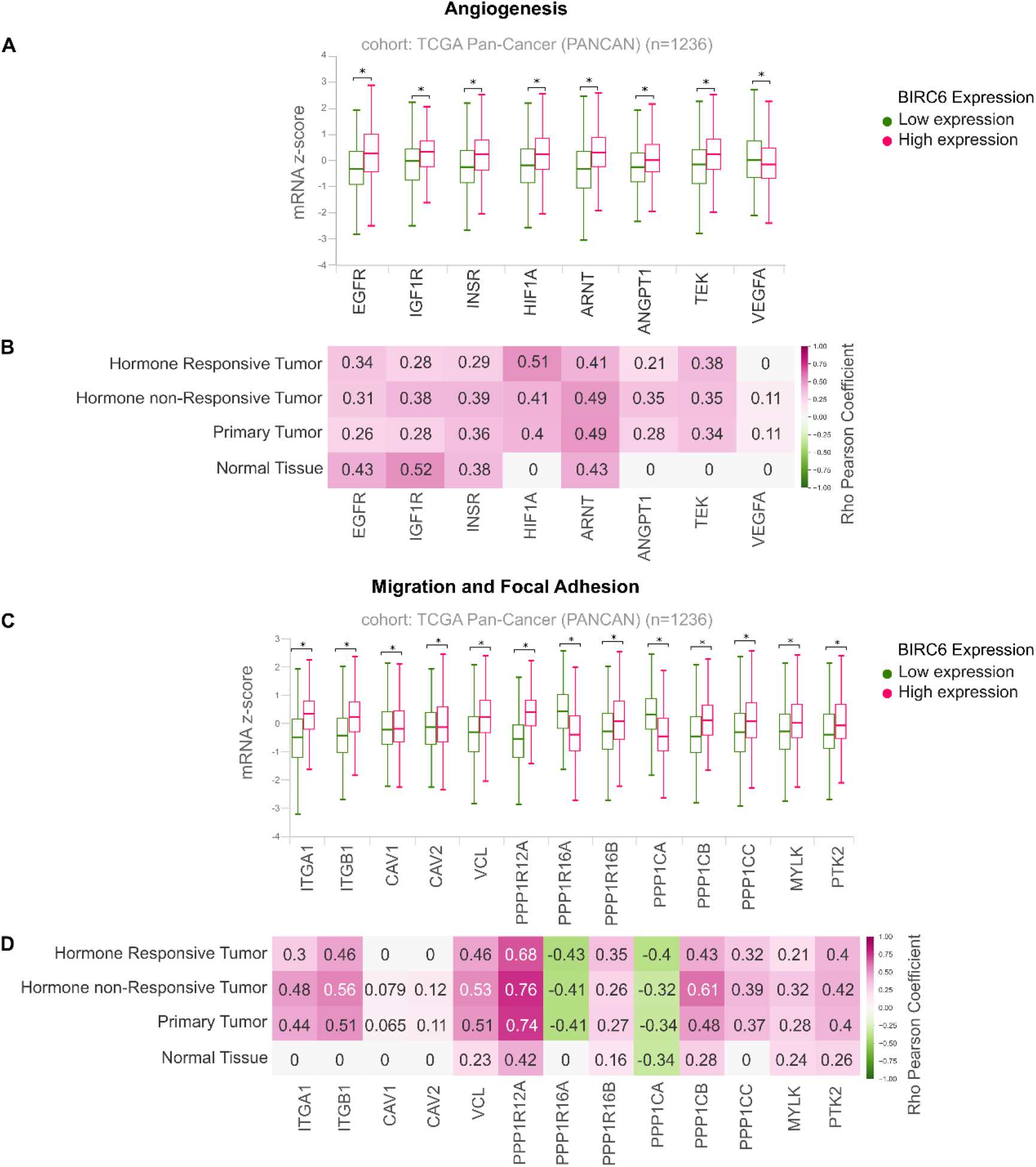
Transcriptomic analysis of pathways related to angiogenesis, migration and focal adhesion (TCGAPanCancer). (A) Boxplot Of The Expression (z-score) of proteins involved in angiogenesis in conditions of high and low expression of BIRC6. T-test *p-value<0.0. (B) Correlation between BIRC6 expression and proteins involved in the angiogenesis pathway separated in normal tissue, primary tumor, hormone responsive and hormone non responsive tumors. Numbers in cells indicate correlation pearson index in statistically significant comparisons. (C) Boxplot of the expression (z-score) of proteins involved in migration and focal adhesion related pathways in condition of high expression and low expression of BIRC6. T-test *p-value<0.01. (D) Correlation between BIRC6 expression and proteins involved in migration and focal adhesion related pathways separated in normal tissue, primary tumor, hormone responsive and hormone non responsive tumors. Numbers in cells indicate correlation pearson index in statistically significant comparisons.

We also evaluated the role of BIRC6 on migration and adhesion processes, since both are necessary for tumor cells to invade other tissues and colonize distant sites, thus generating metastatic nodules. We evaluated the expression of ITGA1, ITGB1, CAV1, CAV2, VCL, PPP1R12A, PPP1R16A, PPP1R16B, PPP1CA, PPP1CB, PPP1CC, MYLK and PTK2 under conditions of low and high BIRC6 expression (Fig 5C). We found that those samples with higher BIRC6 expression presented higher expression of some of the proteins that promote migration such as VCL, PPP1R12A, PPP1CB, MYLK and of those involved in focal adhesions like PTK2.

In addition, we see a distinctive behavior regarding the correlation of BIRC6 and the different proteins analyzed in tumor samples when compared to normal tissue, showing an increased correlation (Fig 5D). This implies a relationship between the expression of BIRC6 and different proteins involved in metastatic cascade. The complete study is presented in S2 Fig and S1 Table.

## 4. Discussion

Breast Cancer is the second leading cause of cancer death worldwide. With the emergence of resistance to conventional therapies arises a growing trend towards the design of new treatment schemes that employ specific molecular targets. Taking this into account, we set out to evaluate the role of BIRC6 in BC development and metastasis by genomic and transcriptomic approaches.

To begin the study, we analyzed BIRC6 expression in different tissues, and we found higher expression levels in samples from mammary carcinomas compared to those corresponding to healthy tissue. This could imply that an increase in the expression of BIRC6 could aid cells in evading the physiological mechanisms of homeostasis, and subsequently become carcinogenic (7,9).

We then assessed the correlation between different clinical attributes and the expression levels of BIRC6 using databases that contained transcriptomic and genomic information of patient samples.

The genomic analysis indicates a higher proportion of amplifications and gains in the number of copies of BIRC6 in ER-and PR-samples when compared to ER+ and PR+. The loss of hormone receptors results in tumors displaying much more aggressive phenotypes (46–51), thus suggesting a possible relationship between BIRC6 expression levels and tumor aggressiveness. We also compared the CNVs with the age of diagnosis and prognosis and found that patients with amplifications or gains in BIRC6 copy number had a tendency to be diagnosed at a younger age. Various studies have shown a correlation between BIRC6 expression and the patient’s prognosis, as well as an involvement of BIRC6 in early stages of cancer development, for colorectal (6), prostate (7,8), and ovarian cancer (9), among others. In the light of this, the relationship of BIRC6 CNV and neoplastic histological grade and the Nottingham prognostic index were analyzed (52–54). We observed that those samples with amplifications or gains showed a tendency to have worse prognosis and to be in more advanced neoplastic degrees. This, together with the loss of the receptors, suggests that an increase in the number of BIRC6 copies could lead to early development and the generation of a more aggressive BC phenotype. To conclude the genomic part of the study, we decided to evaluate the CNV profile in patients who had received either chemotherapy or radiotherapy. It was reported for other types of cancer that BIRC6 could participate in the mechanisms of resistance to these therapies (24,27). We found that in samples of patients who had received chemotherapy or radiotherapy there was a higher proportion of amplifications and gains in gene copy number. Since many signaling pathways are affected in tumor physiology, transcriptomic analysis of the main pathways involved were performed.

We evaluated the state of the different pathways under conditions of high and low BIRC6 expression and the correlation with the different proteins involved, in samples of hormone dependent and independent mammary carcinomas. Regarding apoptosis, gene selection was based on the fact that BIRC6 interacts with different caspases and SMAC/DIABLO, suggesting some kind of correlation in their expression, and also, regulating other points of the pathway that are not directly related. Furthermore, Jinyu Ren et al. reported that the BIRC6 gene is capable of regulating the P53 protein and the mitochondrial apoptotic pathway (15). We observed that the expression of anti-apoptotic genes such as BCL-2 and NFκB1 was increased in high BIRC6 expression samples and pro-apoptotic genes such as BAX, CASP9 and DIABLO were decreased. Despite some effectors with pro-apoptotic activity, like CASP3 and CASP8, correlated with BIRC6 expression, it is important to take into account that BIRC6 interacts directly with both of them exhibiting an inhibitory effect where the net balance results in apoptosis inhibition. Interestingly, despite previous reports in which a relationship between BIRC6 and TP53 was established in breast cancer (22), we found no differences in our transcriptomic analysis.

Another mechanism involved in tumor formation is the unregulated and excessive cell proliferation. It was demonstrated that BIRC6 inhibition decreases tumor cell growth in lung and prostate cancer (8,24). In addition, our results demonstrate that BIRC6 overexpression may promote proliferative pathways like STAT3, PI3KCA, MAPKs and APC in BC.

Tumor metastasis is a multistage process during which malignant cells spread from the primary tumor to non-contiguous organs. The steps that lead to metastasis can be summarized in a few events that are known as the metastatic cascade (55). Considering that metastatic spread affects the survival and prognosis of cancer patients, each of the events that are part of the metastatic cascade are attractive targets for developing therapeutic strategies in oncology. In this work we studied the correlation between BIRC6 expression and key regulators of the metastatic cascade. We observed BIRC6 overexpression correlated with high expression of different proteins involved in migration (such as MYLK), angiogenesis (HIF1A and VEGFA) and adhesion (PTK). Elevated levels of BIRC6 have been linked to cell growth and to some key steps in metastasis in different types of tumors. In lung cancer, BIRC6 overexpression was associated with tumor progression, cell growth, colony formation, migration and invasion as well as with patient metastasis stage (3). In addition, in prostate cancer, the reduction of BIRC6 expression decreases cancer cell viability and proliferation (7,8). We propose that BIRC6 could be involved in tumor progression and metastatic cascade in BC. However, further experiments are needed to explore our proposal *in vivo*.

## 5. Conclusion

In this work, we proposed an integrative study based on genomic and transcriptomic analyses, with the aim of characterizing the role of the apoptosis inhibitor BIRC6 in Breast Cancer. Genomic results show that a higher copy number of BIRC6 gene correlates with more aggressive phenotypes and worse prognosis markers. Transcriptomic studies show that BIRC6 is overexpressed in tumor cells and these levels strongly correlate with an antiapoptotic / proliferative profile. Finally, BIRC6 could play a role in the activation of signaling pathways involved in metastasis.

Altogether our collected data suggest that BIRC6 plays an important role in tumor homeostasis and can be related with the different stages of the metastatic cascade such as angiogenesis, migration and adhesion in BC (Fig 6).

**Fig 6.**
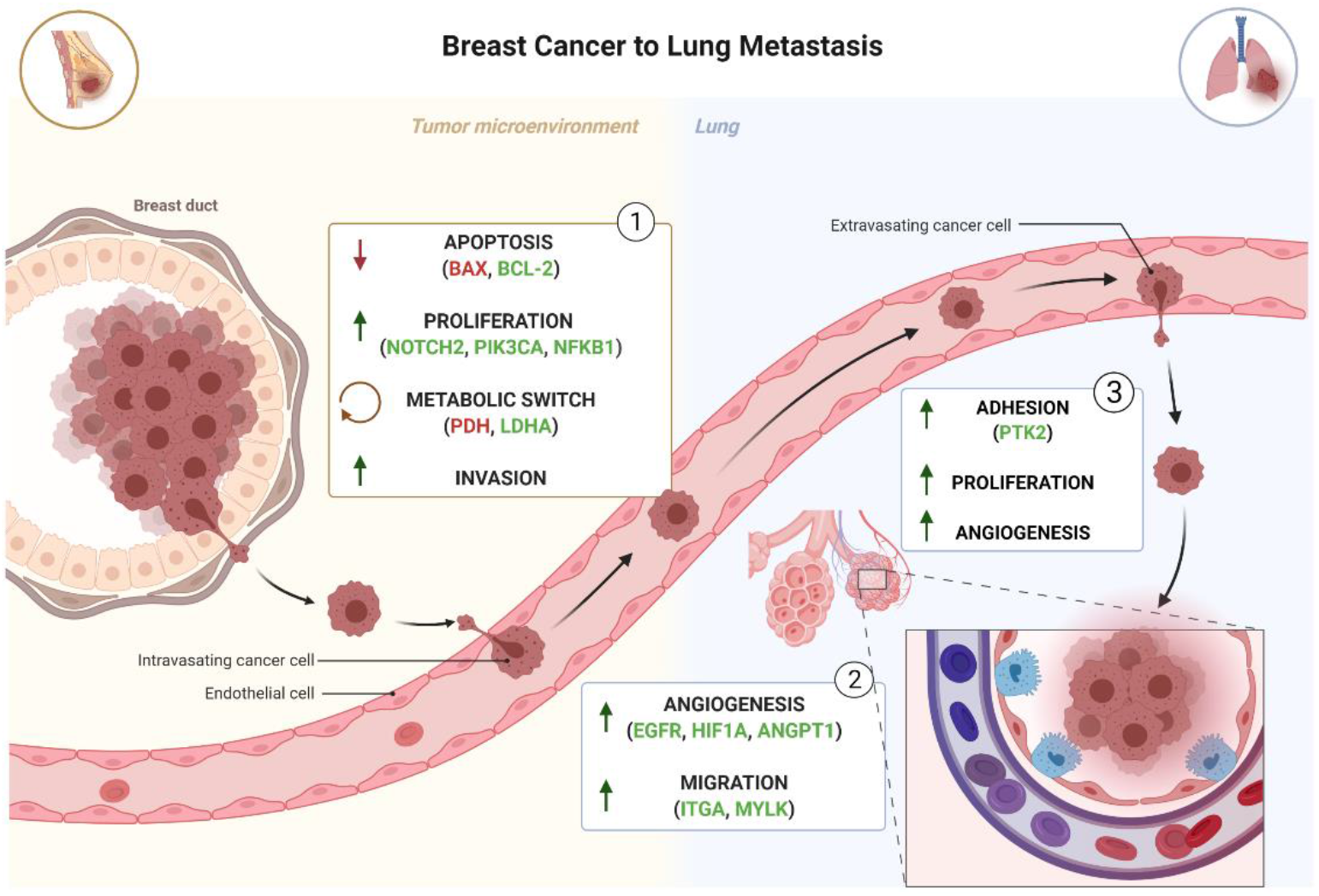
Schematic representation of metastatic cascade. Steps of metastasis are represented in three groups: principal pathways in tumor microenvironment (1), pathways involved in tumor cell spreading (2) and principal pathways in metastatic target organ (3). Principal genes with BIRC6 positive correlation are indicated in green whilst negatively correlated are indicated in red. (Created using BioRender.com).

The results of *in silico* analyses presented here make it possible to postulate the inhibitor of apoptosis BIRC6 as an interesting molecular target for the development of specific therapies for the reduction of tumor progression and metastasis in patients with Breast Cancer.

## Acknowledgements

We thank the *Agencia Nacional de Promoción Científica y Tecnológica* (ANPCyT), the *Consejo Nacional de Investigaciones Científicas y Técnicas* (CONICET), National University of La Plata (UNLP) and National University of Quilmes (UNQ).

