## Supplemental Data for "Exploring the metastatic role of the inhibitor of apoptosis BIRC6 in Breast Cancer"

### Supporting information

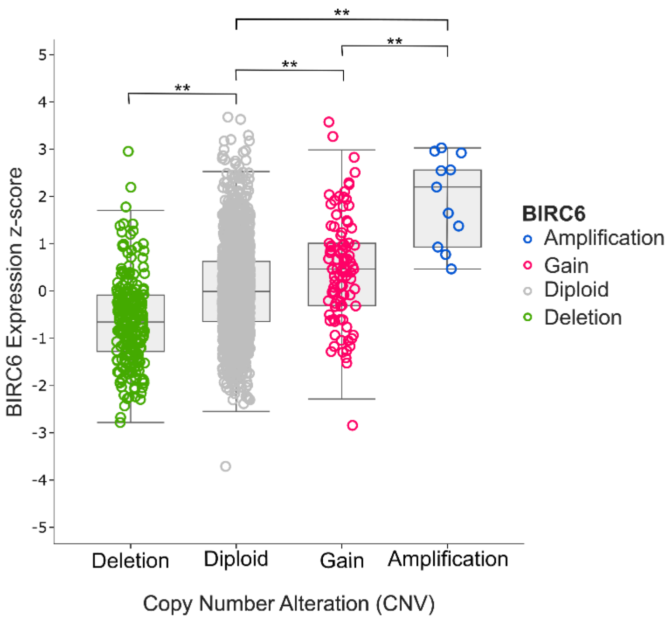


**S1 Fig. BIRC6 expression vs. Copy Number Variation (CNV).** Boxplot of BIRC6 expression z-score in different copy number alterations of birc6. Multiple ANOVA followed by Tukey HSD **p-value<0.001.

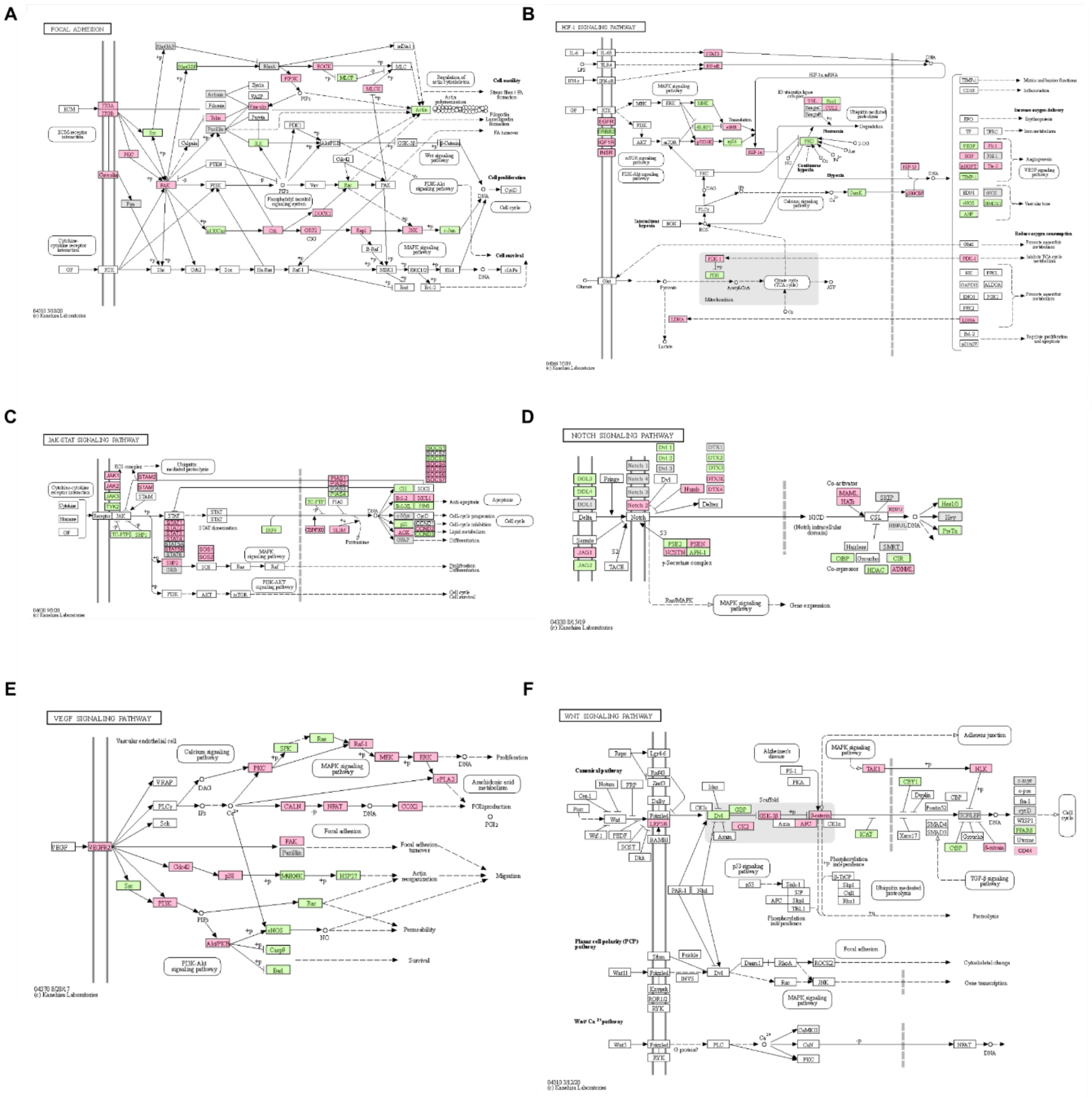


**S2 Fig. Schematic representation of all the pathways analyzed in this work.** (A) Focal Adhesion pathway, (B) HIF1 signaling pathway, (C) JAK-STAT signaling pathway, (D) NOTCH signaling pathway, (E) VEGF signaling pathway and (F) WNT signaling pathway. Genes overexpressed in samples with high levels of BIRC6 are indicated in pink while genes less expressed are indicated in green.

| Focal Adhessions Pathway | | |
|---|---|---|
| Gene | Difference | p-value |
| ITGA1 | UP | 1,86E-49 |
| ITGB1 | UP | 2,97E-31 |
| CAV1 | UP | 0,002735 |
| CAV2 | UP | 0,002591 |
| PRKCA | UP | 5,91E-16 |
| SRC | DOWN | 1,41E-10 |
| PTK2 | UP | 7,25E-11 |
| FYN | NON | 0,5254 |
| ARHGAP1 | NON | 0,5624 |
| ARHGEF1 | DOWN | 0 |
| BCAR1 | DOWN | 0 |
| CRK | UP | 1,05E-61 |
| RASGRF2 | UP | 1,86E-21 |
| RAP1A | UP | 0,000004643 |
| JUN | DOWN | 6,93E-07 |
| MAPK8 | UP | 1,58E-47 |
| RHOA | NON | 0,5136 |
| PIP5K1A | UP | 1,01E-10 |
| PXN | NON | 0,7104 |
| VCL | UP | 1,44E-30 |
| TLN1 | UP | 4,64E-17 |
| ILK | DOWN | 0 |
| ROCK1 | UP | 5,20E-103 |
| PPP1R16A | DOWN | 0 |
| MYLK | UP | 2,25E-13 |
| ACTN1 | NON | 0,5117 |
| ACTN2 | NON | 0,2379 |
| DOCK1 | UP | 5,66E-29 |
| RAC1 | DOWN | 6,10E-09 |
| ACTB | DOWN | 0 |

| JAK/STAT Pathway | | |
|---|---|---|
| Gene | Difference | p-value |
| JAK1 | UP | 1,61E-54 |
| JAK2 | UP | 9,39E-33 |
| JAK3 | DOWN | 0,01867 |
| TYK2 | DOWN | 0 |
| PTPN2 | DOWN | 0,00276 |
| PTPN6 | DOWN | 0 |
| STAM2 | UP | 3,45E-68 |

| Hif-1 alfa Pathway | | |
|---|---|---|
| Gene | Difference | p-value |
| STAT3 | UP | 8,56E-23 |
| NFKB1 | UP | 9,31E-25 |
| HIF1A | UP | 1,90E-16 |
| EGFR | UP | 1,13E-23 |
| ERBB2 | NON | 0,5168 |
| IGF1R | UP | 5,64E-11 |
| INSR | UP | 3,76E-15 |
| MKNK2 | DOWN | 0,0001687 |
| EIF4E | UP | 2,48E-13 |
| EIF4EBP1 | DOWN | 0 |
| RPS6KB1 | UP | 6,89E-24 |
| RPS6 | DOWN | 0 |
| VHL | UP | 8,73E-47 |
| RBX1 | DOWN | 0 |
| CUL2 | UP | 7,08E-28 |
| EGLN2 | DOWN | 0 |
| ARNT | UP | 1,59E-29 |
| CAMK2A | DOWN | 0,0005629 |
| CREBBP | UP | 2,69E-55 |
| VEGFA | DOWN | 0,004588 |
| SLC2A1 | NON | 0,06912 |
| PDK1 | UP | 4,65E-12 |
| PDHA1 | DOWN | 4,73E-13 |
| LDHAL6A | UP | 1,25E-36 |
| FLT1 | UP | 4,39E-22 |
| SERPINE1 | NON | 0,9315 |
| EGF | UP | 4,67E-07 |
| ANGPT1 | UP | 5,76E-13 |
| TEK | UP | 3,72E-19 |
| TIMP1 | DOWN | 0 |
| NOS2 | NON | 0,6382 |
| HMOX1 | DOWN | 7,67E-08 |
| NOS3 | DOWN | 0,0000163 |
| NPPA | DOWN | 0,0009015 |

| NOTCH Pathway | | |
|---|---|---|
| Gene | Difference | p-value |
| NOTCH1 | NON | 0,5009 |
| NOTCH2 | UP | 4,64E-81 |
| NOTCH3 | NON | 0,5132 |

| | | |
|---|---|---|
| STAM | UP | 1,15E-30 |
| STAT1 | UP | 3,37E-09 |
| STAT2 | UP | 0,000001381 |
| STAT3 | UP | 8,56E-23 |
| STAT4 | UP | 0,01146 |
| STAT5A | NON | 0,7322 |
| STAT5B | UP | 1,07E-08 |
| STAT6 | NON | 0,6232 |
| IRF9 | DOWN | 7,55E-11 |
| PIAS3 | NON | 0,8945 |
| PIAS4 | DOWN | 0,000006154 |
| PIAS1 | UP | 4,77E-102 |
| PIAS2 | UP | 3,73E-35 |
| CREBBP | UP | 2,69E-55 |
| EP300 | UP | 3,37E-88 |
| FHL1 | UP | 0,000004856 |
| CISH | DOWN | 0,0000209 |
| BCL2 | UP | 2,34E-11 |
| BCL2L1 | DOWN | 2,25E-08 |
| PIM1 | DOWN | 0,0006126 |
| MCL1 | UP | 0,00005153 |
| SOCS1 | DOWN | 0 |
| SOCS2 | NON | 0,09893 |
| SOCS3 | DOWN | 0,002348 |
| SOCS4 | UP | 1,41E-63 |
| SOCS5 | UP | 2,35E-48 |
| SOCS6 | UP | 1,48E-34 |
| SOCS7 | UP | 3,12E-84 |
| MYC | NON | 0,1119 |
| CCND1 | NON | 0,09984 |
| CCND2 | UP | 0,00003168 |
| CCND3 | DOWN | 0 |
| CDKN1A | DOWN | 0,01158 |
| AOX1 | UP | 3,59E-12 |
| GFAP | NON | 0,8869 |
| PTPN11 | UP | 1,96E-83 |
| GRB2 | NON | 0,9086 |
| SOS1 | UP | 1,64E-90 |
| SOS2 | UP | 6,09E-68 |

**VEGF Pathway**

| Gene | Difference | p-value |
|---|---|---|

| | | |
|---|---|---|
| NOTCH4 | NON | 7,10E-01 |
| DLL3 | DOWN | 2,39E-05 |
| DLL1 | NON | 0,8671 |
| DLL4 | DOWN | 2,22E-04 |
| JAG1 | UP | 5,80E-07 |
| JAG2 | DOWN | 4,66E-15 |
| DVL1 | DOWN | 0,00E+00 |
| DVL2 | DOWN | 2,44E-13 |
| DVL3 | NON | 7,51E-01 |
| NUMB | UP | 3,37E-27 |
| DTX2 | DOWN | 0 |
| DTX1 | NON | 3,28E-01 |
| DTX3 | DOWN | 1,33E-15 |
| DTX3L | UP | 5,53E-16 |
| DTX4 | UP | 0,01429 |
| PSENEN | DOWN | 0 |
| PSEN1 | UP | 1,19E-10 |
| NCSTN | UP | 2,34E-03 |
| APH1A | DOWN | 0 |
| RBPJ | UP | 1,76E-20 |
| RBPJL | NON | 0,7691 |
| MAML3 | UP | 1,77E-19 |
| CREBBP | UP | 2,69E-55 |
| SNW1 | NON | 0,9123 |
| CTBP1 | DOWN | 0 |
| HDAC1 | DOWN | 0,000001714 |
| ATXN1L | UP | 5,79E-74 |
| CIR1 | DOWN | 1,47E-08 |
| NCOR2 | NON | 0,5471 |
| HES1 | DOWN | 0 |
| HES5 | DOWN | 1,03E-08 |
| HEY1 | NON | 0,3193 |
| HEY2 | NON | 0,8819 |
| HEYL | NON | 0,9933 |
| PTCRA | DOWN | 2,98E-10 |

**Wnt/beta-catenin Pathway**

| Gene | Difference | p-value |
|---|---|---|
| FZD1 | NON | 7,14E-02 |
| LRP6 | UP | 1,23E-52 |
| CTNNB1 | UP | 1,25E-31 |
| APC | UP | 2,08E-113 |

| | | | | | |
|---|---|---|---|---|---|
| KDR | UP | 5,18E-22 | GSK3B | UP | 6,30E-74 |
| PLCG1 | NON | 5,94E-02 | FRAT1 | DOWN | 5,37E-12 |
| PRKCA | UP | 5,91E-16 | CTNNBIP1 | DOWN | 6,44E-15 |
| SPHK2 | DOWN | 0,00E+00 | CBY1 | DOWN | 0,00E+00 |
| HRAS | DOWN | 0 | NLK | UP | 3,58E-04 |
| RAF1 | UP | 6,82E-06 | MAP3K7 | UP | 3,76E-40 |
| MAP2K1 | UP | 6,79E-07 | TAB1 | DOWN | 0,00E+00 |
| MAPK1 | UP | 3,61E-61 | TAB2 | UP | 1,30E-48 |
| PLA2G4A | UP | 9,66E-08 | MYC | NON | 0,1119 |
| PPP3CA | UP | 2,93E-12 | CCND1 | NON | 9,98E-02 |
| NFATC2 | UP | 1,32E-67 | PPARD | DOWN | 0,000001985 |
| PTGS2 | UP | 0,0003713 | MMP7 | NON | 5,90E-02 |
| PTK2 | UP | 7,25E-11 | AXIN2 | UP | 3,96E-08 |
| PXN | NON | 0,7104 | AXIN1 | DOWN | 0 |
| CDC42 | UP | 2,26E-03 | CD44 | UP | 3,11E-03 |
| MAPK14 | UP | 1,52E-49 | POU1F1 | NON | 6,50E-02 |
| MAPKAPK3 | DOWN | 1,56E-11 | HDAC1 | DOWN | 1,71E-06 |
| HSPB1 | DOWN | 0 | DVL1 | DOWN | 0,00E+00 |
| SRC | DOWN | 1,41E-10 | LEF1 | NON | 0,994 |
| PIK3CA | UP | 3,48E-79 | CSNK2A1 | UP | 8,81E-17 |
| AKT3 | UP | 3,56E-17 | CTNND2 | UP | 8,71E-03 |
| NOS3 | DOWN | 1,63E-05 | CTBP1 | DOWN | 0,00E+00 |
| CASP9 | DOWN | 6,61E-05 | | | |
| BAD | DOWN | 0,00E+00 | | | |
| RAC1 | DOWN | 6,10E-09 | | | |

**S1 Table. Summary of all the genes analyzed in this work involved in:** Focal Adhesion pathway, HIF1 signaling pathway, JAK-STAT signaling pathway, NOTCH signaling pathway, VEGF signaling pathway and WNT signaling pathway. Genes overexpressed (UP) in samples with high levels of BIRC6 are indicated in pink while genes less expressed (DOWN) are indicated in green. Genes with no changes are indicated in grey (NON). T-test statistical analysis was performed for each gene comparing conditions of high vs. low BIRC6 expression.

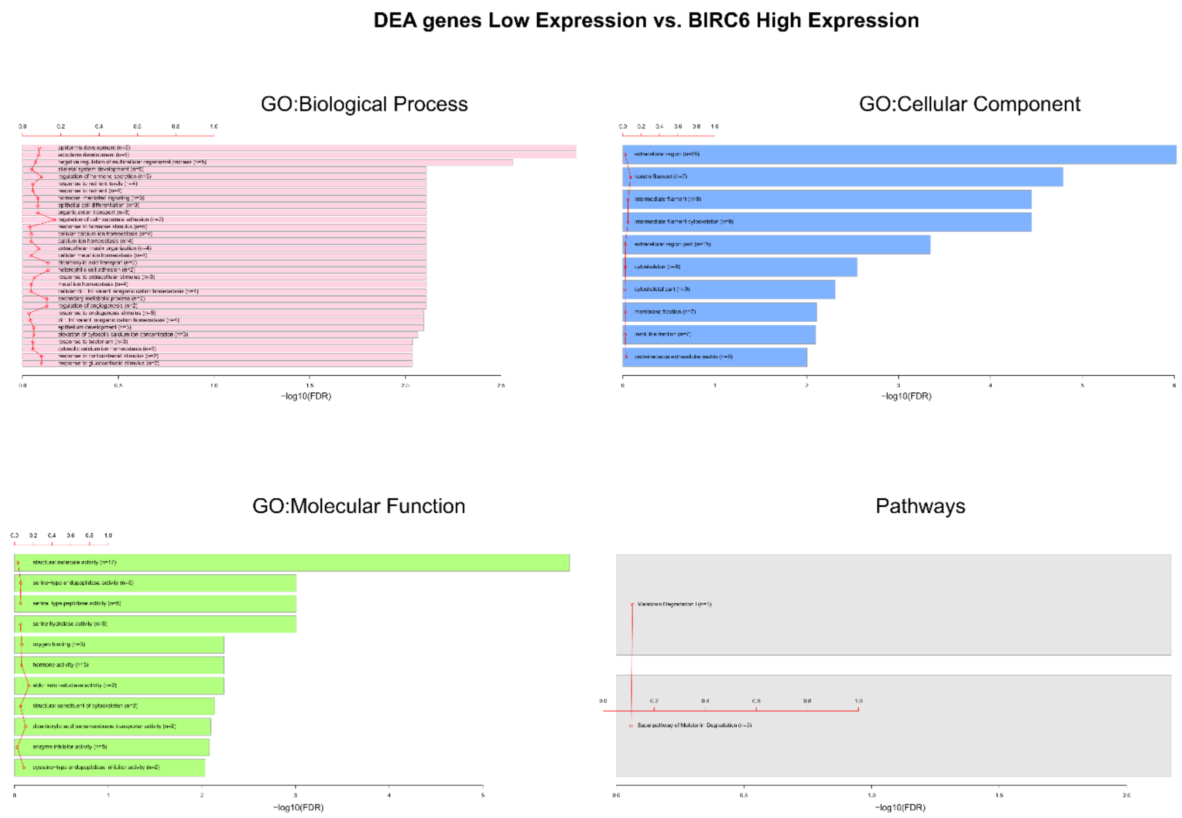


**S3 Fig. Gene ontology and down-represented pathways.** Graphs show the canonical pathways significantly down-represented by the DEGs (differentially expressed genes) with the number of genes for the main categories of the three ontologies (GO: biological process, GO: cellular component, and GO: molecular function, respectively). The statistically significant canonical pathways in the DEGs are listed according to their p-value corrected by FDR (- log10) (colored bars) and the ratio of the listed genes found in each pathway over the total number of genes in that pathway (ratio, red line).
